# TRANSPOSABLE ELEMENTS ALTER GENE EXPRESSION AND MAY IMPACT RESPONSE TO CISPLATIN THERAPY IN OVARIAN CANCER

**DOI:** 10.1101/2023.09.29.560129

**Authors:** Daniela Moreira Mombach, Rafael Luiz Vieira Mercuri, Tiago Minuzzi Freire da Fontoura Gomes, Pedro A. F. Galante, Elgion Lucio Silva Loreto

## Abstract

Cisplatin is widely employed for cancer treatment; therefore, understanding resistance to this drug is critical for therapeutic practice. While studies have delved into differential gene expression in the context of cisplatin resistance, findings remain somewhat scant. In this study, we employed RNA-seq, ATAC-seq, and in-depth bioinformatics analyses to perform a detailed investigation of the cellular transcriptome, centering on Transposable Elements (TEs) expression in ovarian cancer cell lines both sensitive and resistant to cisplatin treatment. Our results reveal that cisplatin therapy alters the expression of protein-coding genes, but also key TEs, including LINE1, *Alu*, and endogenous retroviruses, in both cisplatin-sensitive and -resistant cell lines. By co-expressing with downstream genes or by creating chimeric transcripts with host genes at their insertion sites, these TEs seem to control the expression of protein-coding genes, including tumor-related genes. Notably, our model uncovers TEs influencing the expression of cancer genes and cancer pathways. Collectively, our findings indicate that TEs alterations associated with cisplatin treatment occur in critical cancer genes and cellular pathways synergically. In conclusion, this research highlights the importance of considering the entire spectrum of transcribed elements in the genome, especially TE expression, for a complete understanding of complex models like cancer response to treatment.

## INTRODUCTION

Cisplatin exerts multiple anticancer effects. Its primary and best-understood mechanism of action is the induction of DNA lesions, subsequently triggering the DNA damage response and leading to mitochondrial-induced cell death^1^. However, cisplatin’s effectiveness as a broad-spectrum anticancer drug is hindered by the prevalent issue of chemoresistance. Such resistance can arise through various molecular mechanisms, including increased detoxification system, increased DNA damage repair, and the evasion of apoptosis^2^. While the expression profiles of protein-coding genes have been the subject of numerous studies in cisplatin-resistant and -sensitive ovarian cancer (OC)^3–5^, the findings have often been conflicting or limited.

Transposable elements (TEs) constitute nearly half of the human genome^6^. While commonly recognized for their mobility, TEs can also influence cellular gene expression. De-repression of specific TE loci can impact transcription or processing, modulate gene expression levels, and alter chromatin accessibility of protein-coding genes^7–9^. Reactivation of TEs is reported in a variety of malignancies, and is associated with oncogenic factors such as oncogene activation^10^, genome instability promotion^11^, genomic mutation occurrence^11^, putative immunotherapy roles^12^, and even drug treatment influences^13^.

Furthermore, TEs are a rich source of cis-regulatory elements that drive changes in protein-coding gene expression^14^. The myriad of transcription factor binding sites (TFBSs) originated from TE sequences, coupled with their inherent regulatory features, underscore the diversity of TE-mediated cis-regulation of gene expression^15^. Experimental and genome-wide analytical studies have pinpointed TE sequences in numerous gene regulatory regions, with certain genes demonstrably regulated by TEs^16,17^. The shifts in TE expression and their roles in gene regulation have been documented in various contexts, most notably in cancers^10^, accentuating the urgency for further targeted research in this field.

Despite diagnostic and therapeutic advances, OC remains the fifth leading cause of cancer-related death among women, emerging as the deadliest gynecologic malignancy^18^. In addition to difficulties with early-stage detection, the high mortality rate of OC is also due to the development of multidrug resistance, especially against platinum-based drugs like cisplatin. Cisplatin has been a cornerstone in OC chemotherapy for decades^19^, yet resistance (intrinsic or acquired) poses a substantial therapeutic barrier^2^. This scenario emphasizes the urgency of delving deeper into the molecular mechanisms underpinning cisplatin resistance.

In this study, we carried out an extensive investigation of TEs expression in two OC cell lines: one cisplatin-sensitive and the other cisplatin-resistant, both treated with cisplatin. Our findings provide compelling evidence suggesting TEs play an integral role in regulating critical cancer genes across multiple mechanisms, potentially affecting cisplatin response. This insight sheds light on the potential of TEs as novel therapeutic targets in cancer research.

## RESULTS

### Overview of our study approach

To investigate TEs and gene expression, we designed a straightforward study approach (**Fig. 1**). Initially, we gathered RNA sequencing (RNA-seq) and transposase-accessible chromatin followed by sequencing (ATAC-seq) data from the cisplatin-sensitive A2780 ovarian cancer cell line and its cisplatin-resistant derivative, A2780cis. The gene and TE expression data were subjected to differential expression analysis to determine their deregulation following cisplatin treatment. Subsequently, we employed a multistep pipeline to assess the impact of TEs on the regulation and expression of genes, protein-coding genes, and lncRNAs in this cellular model. To investigate the TEs associated with differentially expressed genes (DEGs), we followed with other approaches, such as TE-gene proximity (intersection) and the formation of genetic novelties (chimeric transcripts). DEGs associated with TEs were searched for in regions that were uniquely accessible (ATAC-seq data) after cisplatin treatment and used in a functional enrichment analysis. Finally, TE enrichment was searched in these new regions after cisplatin treatment (**Fig. 1**). The results also included comparisons between cisplatin-sensitive and -resistant cell lines.

**Figure 1.**
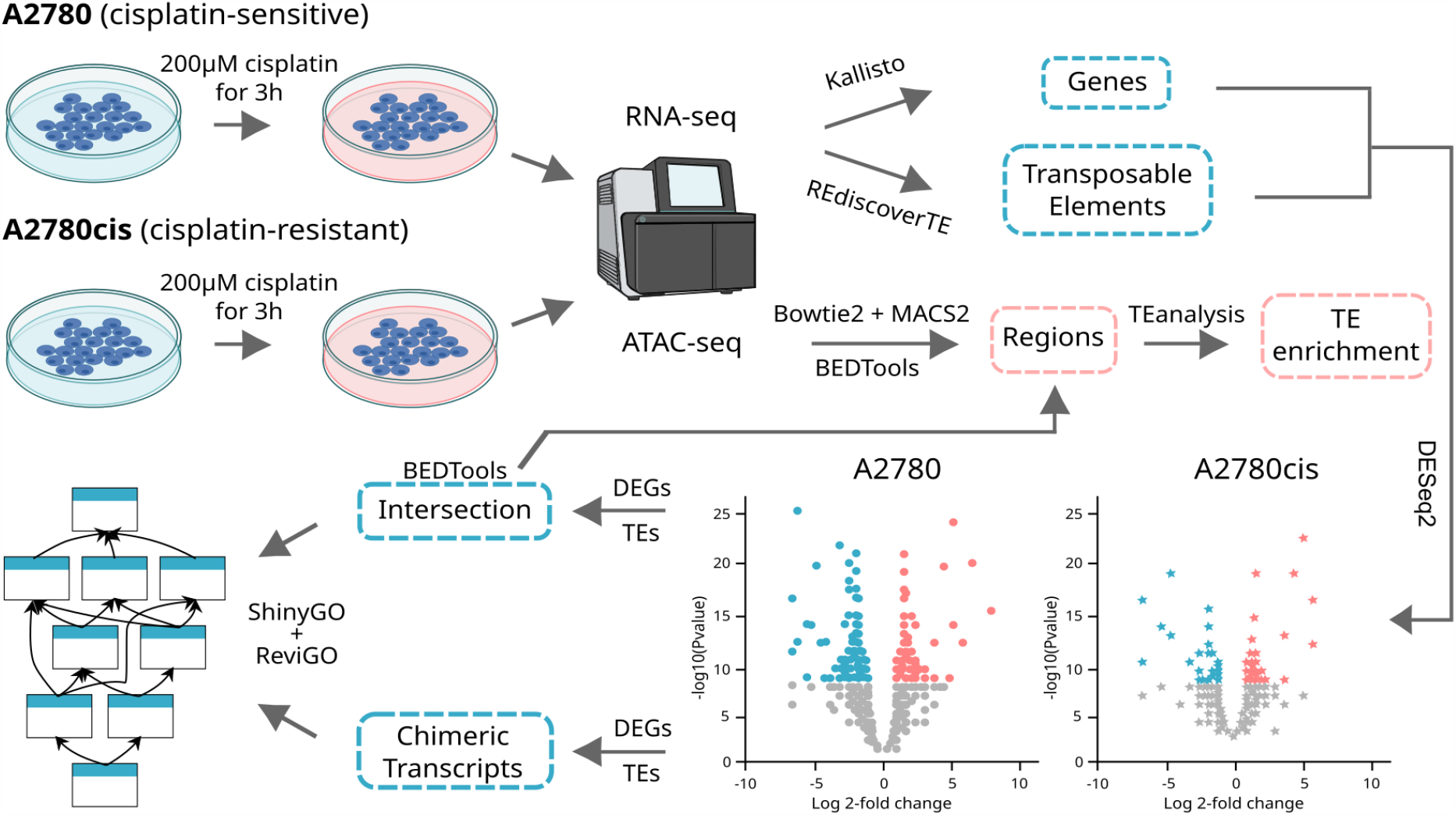
Approach employed to investigate the impact of transposable elements on gene expression in cisplatin-treated ovarian cancer cell lines. RNA-seq data from cisplatin-sensitive and -resistant OC cell lines treated with cisplatin were used to quantify gene and TE expression using Kallisto^20^ and REdiscoverTE^21^, respectively. ATAC-seq data were used to identify chromatin accessibility regions using Bowtie2^22^, MACS2^23^ and BEDtools^24^ intersect. Expression results were then followed by differential expression analysis of both genes and TEs using DESeq2^25^. The interactions of TEs with genes were further investigated via coordinate crossing, using BEDTools intersect, and chimeric transcripts formation using Freddie (Mercuri RLV, Rangel A, and Galante PAF, manuscript in prep.). Differentially expressed genes impacted by TEs were searched for in regions that were uniquely accessible after cisplatin treatment and used to identify functional enrichment using ShinyGO^26^ followed by ReviGO^27^. TE enrichment in these regions was determined using the Perl script. See Methods for detail.

### Cisplatin-induced differential gene expression profiles

Previously to the investigation of the TEs in both cell lines under treatment and control conditions, we decided to profile the coding and non-coding gene expression in these cell lines. First, we sought the set of DEGs in both the A2780 and A2780cis cell lines under treatment and control conditions. For the A2780cis (cisplatin-resistant) cell line we identified 465 DEGs in total (220 protein-coding), of which 266 were up-regulated and 199 were down-regulated after cisplatin treatment. Similarly, for A2780 we found 1 624 DEGs (695 protein-coding), of which 1 000 were up-regulated and 624 were down-regulated (**Fig. 2A** and **Supplementary Table 1**). Next, we assessed the overlap of DEGs between the two cell lines. Intriguingly, we identified a small subset of shared protein-coding genes (44; representing 5.05% of protein-coding DEGs) between A2780 (cisplatin-sensitive) and A2780cis (cisplatin-resistant) cell lines (**Supplementary Table 2**). This suggests that cisplatin treatment either induces or suppresses the transcription of specific genes in each cell line under treatment. **Fig. 2B** displays a heatmap of these protein-coding DEGs, highlighting the unique gene expression patterns observed between cell lines, as well as the differential gene expression profiles within each cell line when comparing control and treatment conditions. Therefore, collectively these results confirm the findings of other authors that show a higher number of DEGs in A2780^4^. Moreover, it also demonstrates that, in addition to there being sets of genes that are differentially expressed due to cisplatin, the two cell lines exhibit entirely distinct profiles of differentially expressed genes when subjected to treatment, in line with the fact that they are sensitive (A2780) or resistant (A2780cis) to cisplatin.

**Figure 2.**
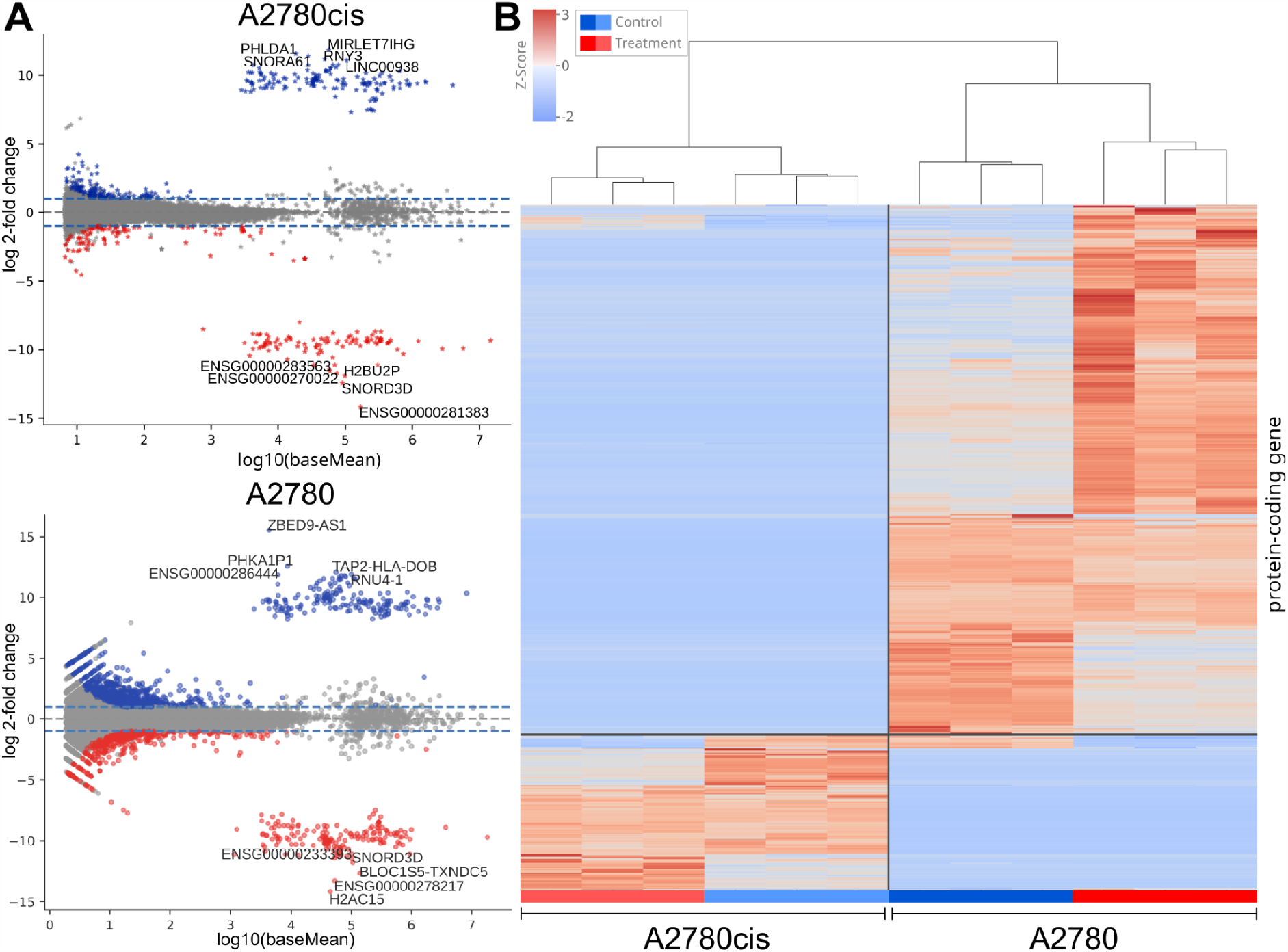
Differentially expressed genes after cisplatin treatment. **A)** MA plot depicting the differentially expressed genes (DEGs). MA plots were color-coded according to the expression of the gene, either up-regulated in blue or down-regulated in red (log 2-fold change lower than -1 or greater 1, and adjusted p-value (False Discovery Rate) below 5%). Grey is for non-significant genes. Dashed lines represent log 2-fold change lower than -1 or greater 1. **B)** Heatmap displaying the expression levels of the protein-coding DEGs, represented by Z-Scores of TPM (Transcripts Per Million). The DEGs were identified based on an adjusted p-value (False Discovery Rate) below 5% and a log 2-fold change lower than -1 or greater 1.

### Cisplatin-induced differential transposable element expression profiles

It is noteworthy that approximately half of the human genome consists of TEs^6^, which are tightly regulated and commonly repressed in differentiated cells. However, in the context of cancer, TEs expression regulation becomes impaired, allowing TEs to be frequently transcribed. Therefore, we decided to explore the transcriptomics of TEs during cisplatin treatment. By examining the set of differentially expressed TEs (DETEs) in both the cisplatin-resistant (A2780cis) and -sensitive (A2780) cell lines under treatment and control conditions, we identified differential expression of LINE1 (L1), *Alu* elements, long terminal repeats (LTRs; referred to as ERVs in this study), and other TE families (e.g., SVAs). For the A2780cis cell line we found 47 DETEs in total, of which 30 TEs were up-regulated and 17 were down-regulated. Similarly, in the A2780 cell line, we found 60 DETEs, of which 33 were up-regulated and 27 were down-regulated (**Fig. 3A** and **Supplementary Table 3**). Consistent with the DEGs analysis, this analysis revealed distinct expression profiles of DETEs associated with the two cell lines under treatment (**Fig. 3B**), and a higher number of elements being differentially expressed in cisplatin-sensitive cell line (A2780). We observed only three shared DETEs across cell lines (**Supplementary Table 4**), including an intriguing TE, PrimLTR79 (an ERV1 subfamily member), which was up-regulated in the cisplatin-resistant cell line, but down-regulated in A2780. When comparing the DETEs families between cell lines, ERVs emerged as the dominant component, constituting 76.6% of DETEs in A2780cis and 63.3% in A2780 (**Fig. 3C**). Within the broad ERV category, the ERV1 and ERVL subfamilies were the most prevalent in the DETEs of both cell lines (**Fig. 3C**). Overall, our findings indicate that cisplatin treatment can either induce or repress TE expression, and the specific TEs expressed differ between cisplatin-sensitive and -resistant cell lines. Notably, among DETEs, ERVs stand out as the most abundant, emphasizing their potential role in the cellular response to cisplatin treatment.

**Figure 3.**
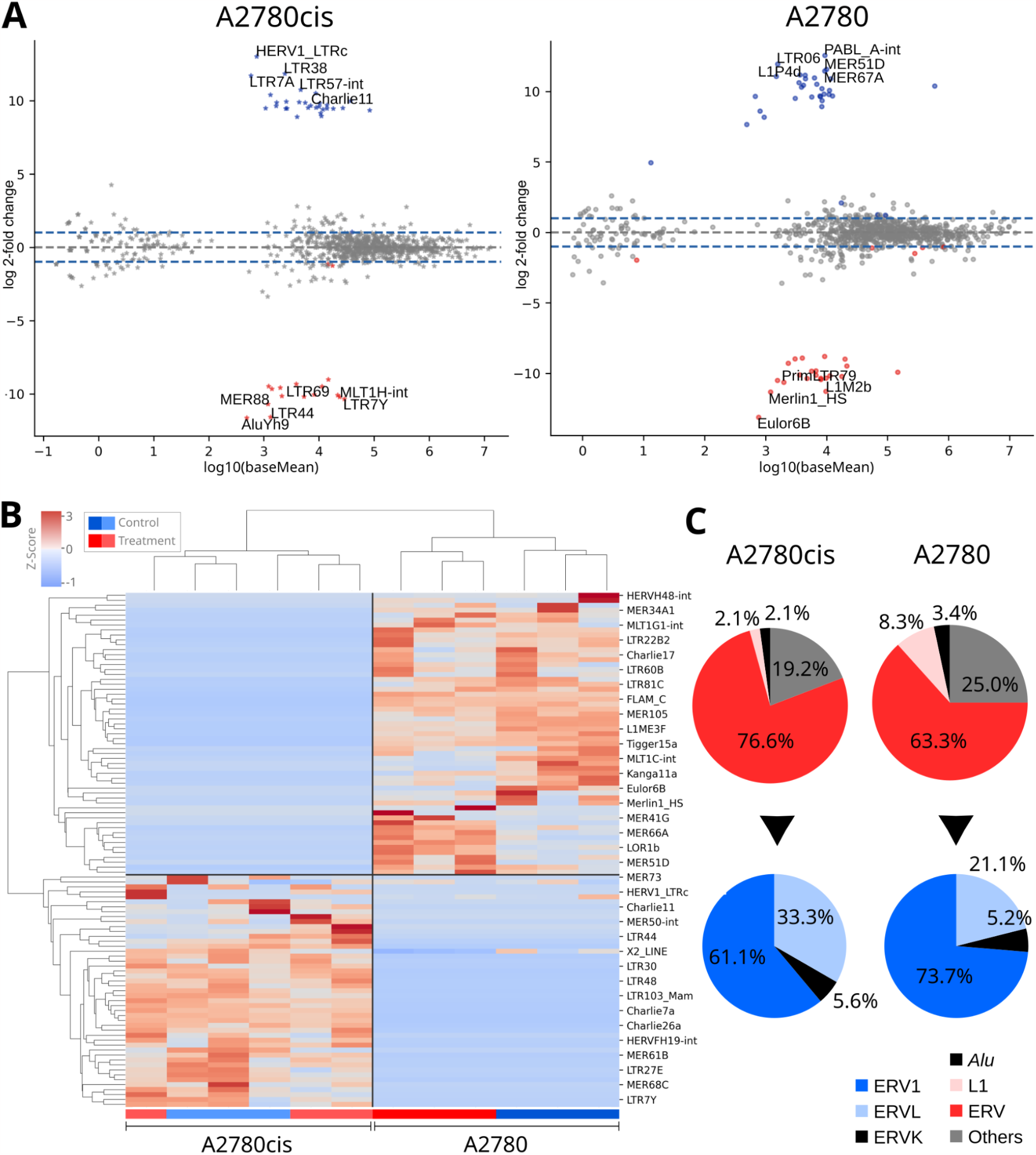
Differentially expressed transposable elements after cisplatin treatment. **A)** MA plot depicting the differentially expressed transposable elements (DETEs). MA plots were color-coded according to the expression of the TE, either up-regulated in blue or down-regulated in red (log 2-fold change lower than -1 or greater 1, and adjusted p-value (False Discovery Rate) below 5%). Grey is for non-significant TEs. Dashed lines represent log 2-fold change lower than -1 or greater 1. **B)** Heatmap of DETEs using Z-Score of TPM. **C)** DETEs first grouped into TE families, and then into subfamilies of endogenous retroviruses. The DETEs were identified based on an adjusted p-value (False Discovery Rate) below 5% and a log 2-fold change lower than -1 or greater 1.

### Transposable elements proximity to differentially expressed cancer-related genes

Given that cisplatin-sensitive and -resistance cells present a distinct set of DEGs and DETEs, including TEs with regulatory functions^9^, we next asked whether transcribed TEs are cis-associated with DEGs (**Fig. 4A-B**). First, using DEGs and TEs from each respective cell line and condition, we found transcribed TEs within promoter regions (up to 5 Kb) or inside DEGs (**Supplementary Table 5**). In accordance with the number of DEGs and DETEs, A2780 (treatment versus control condition) presented roughly three times more TEs upstream the promoter region of DEGs (**Fig. 4A**) and inside the transcribed region of DEGs than A2780cis (**Fig. 4B**; **Supplementary Table 5**). Interestingly, we observed distinct TE profiles between those upstream and inside up-regulated DEGs (**Fig. 4C**). While TEs upstream (near or inside promoter region) are enriched to ERVs and L1, TEs inside genes are enriched to *Alu* elements (**Fig. 4C**).

**Figure 4.**
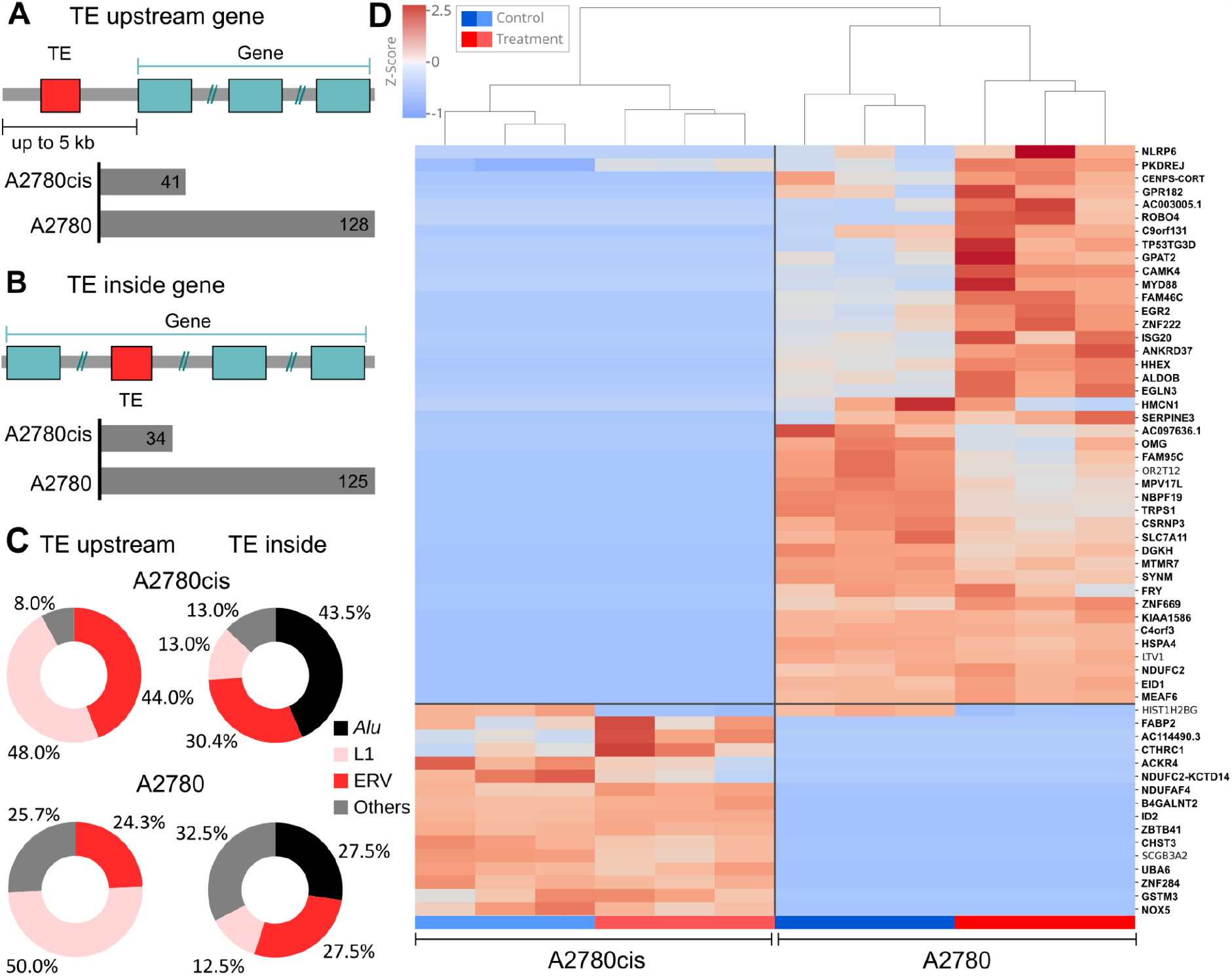
Transposable elements coordinates were crossed with the genes for two potential outcomes. **A)** TEs upstream differentially expressed genes (DEGs), with up to 5 kb from the gene’s promoter region, or **B)** TEs inside the DEGs’ region. **C)** The profile of TE upstream and inside up-regulated DEGs grouped into TE family. **D)** Clustered heatmap of differentially expressed protein-coding genes with TEs either inside or upstream their sequences. DEGs with TE upstream were highlighted in bold. DEGs were identified based on an adjusted p-value (False Discovery Rate) below 5% and a log 2-fold change lower than -1 or greater 1. TEs inside or upstream DEGs were filtered for TPM higher than 1.

To further observe the expression profile of these protein-coding DEGs with TEs inside and upstream, we proceeded with a clustered heatmap analysis (**Fig. 4D**; **Supplementary Fig. 1**). We observed within cell lines those genes associated with the treatment, but the difference between the two cells considering these genes is remarkably clear. By picking some genes and their associated TEs, we highlight a L1M3 and an ERVK (LTR5B) family members present upstream the sequences of the up-regulated tumor suppressor gene *FAM46C* and the up-regulated oncogene *MYD88*, respectively (both up-regulated in A2780 cell line; **Supplementary Fig. 2**). Together, these findings suggest a cis-association between differentially expressed genes and TEs, hinting at a regulatory role for the latter. Intriguingly, the patterns of differential expression for both genes and associated TEs unequivocally distinguish between cisplatin-sensitive and -resistant cell lines. Finally, some TEs are located near or within genes tied to carcinogenesis, potentially influencing the response to cisplatin treatment.

### Regulatory mechanisms for differentially expressed genes and transposable elements in response to cisplatin treatment

To gain more insight about TEs regulating gene expression in our model, we evaluated changes in DNA accessibility associated with DEGs and TEs after cisplatin treatment with ATAC-seq. By analyzing ATAC-seq data from both cell lines, A2780 and A2780cis, under cisplatin treatment and control, we found 6 827 and 7 283 peaks uniquely accessible after cisplatin treatment in A2780 and A2780cis, respectively. Next, crossing these peaks with nearby DEGs (+1 kb upstream their TSS) up-regulated in A2780 and A2780cis, we found 141 and 47 differentially expressed genes, respectively (**Supplementary Table 6**). We next explored the functional enrichment of these genes using the Gene Ontology (GO) biological process classification. Consistent with cisplatin sensitivity, the A2780 cell line exhibited enrichment in biological processes related to the regulation of apoptotic processes and cell death, as well as the regulation of the cell cycle and cellular responses to DNA damage (**Supplementary Table 7**; **Supplementary Fig. 3**). Due to the limited number of genes (47), the A2780cis showed no enrichment in this analysis. Finally, we focused on up-regulated DEGs with expressed TEs either inside or upstream their sequences, and we found 11 and 3 genes for A2780 and A2780cis, respectively (**Supplementary Table 8**). We next intersected the ATAC-seq peaks with TE coordinates (±1 kb). We searched if the TE families inside or upstream up-regulated DEGs were not only enriched in the new regions, but also up-regulated after cisplatin treatment, in order to identify specific TE families undergoing chromatin changes and subsequent transcriptional activation. Only one up-regulated TE (SVA family), was enriched in newly accessible chromatin regions after cisplatin treatment in A2780 cell line (**Supplementary Table 9**). In summary, the chromatin accessibility assays revealed associations with both DEGs and expressed TEs in both cell lines, suggesting distinct regulatory mechanisms for genes (and potentially TEs) in response to cisplatin treatment, particularly in the A2780 cell line.

### Cisplatin induces chimeras of cancer-related genes and transposable elements

Based on our previous results (**Fig. 3-4**), and given that TEs inserted within or adjacent to gene sequences can profoundly influence host gene expression, we explored the potential formation of chimeric transcripts between TEs and their host genes, as illustrated in **Fig. 5A**. This analysis revealed 17 chimeric transcripts in the A2780 cell line (16 associated with *Alu* elements and one with L1), and one L1 chimeric transcript in the A2780cis cell line (**Supplementary Table 10**). Several intriguing candidates emerged. For instance, we identified a chimeric transcript (resulting from *AluY* exonization) in the *CDKN1A* gene (also known as p21). *CDKN1A* is a pivotal regulator of the cell cycle, and consistent with expectations, it is up-regulated in A2780 following treatment. The chimeric transcript retains the *CDKN1A* ORF (**Fig. 5B**), leading us to hypothesize that the chimeric version, in partnership with TEs, might enhance the gene’s activity, thereby promoting cell death when treated. Conversely, we detected a chimeric transcript associated with the *DDB2* gene that generates a non-functional transcript due to a premature stop codon (**Fig. 5C**). Given that *DDB2* (Damage-Specific DNA Binding Protein 2) encodes a protein integral to the nucleotide excision repair pathway (a primary mechanism cells utilize to mend DNA damage) the presence of this non-functional chimeric transcript could influence the responsiveness of A2780 cells to cisplatin treatment. Another notable chimeric transcript, which does not code for a functional protein (**Fig. 5D**), was found in the *BCL2L12* gene. *BCL2L12* (BCL2 Like 12) has a significant role in regulating apoptosis. It is typically characterized as an anti-apoptotic protein, and elevated expression levels of *BCL2L12* have been documented in various cancers, suggesting its potential involvement in tumor survival and therapy resistance. Therefore, discovering a nonfunctional chimeric transcript in this gene hints at a heightened response to cisplatin treatment, consistent with observations in A2780. Details on all other chimeric transcripts are provided in **Supplementary Table 10**. Collectively, these findings demonstrate that TEs embedded in our genome (either within or proximal to genes) can form chimeric transcripts with host genes in response to specific treatments, such as cisplatin. Here, the genes impacted align with the cell’s sensitivity to the treatment.

**Figure 5.**
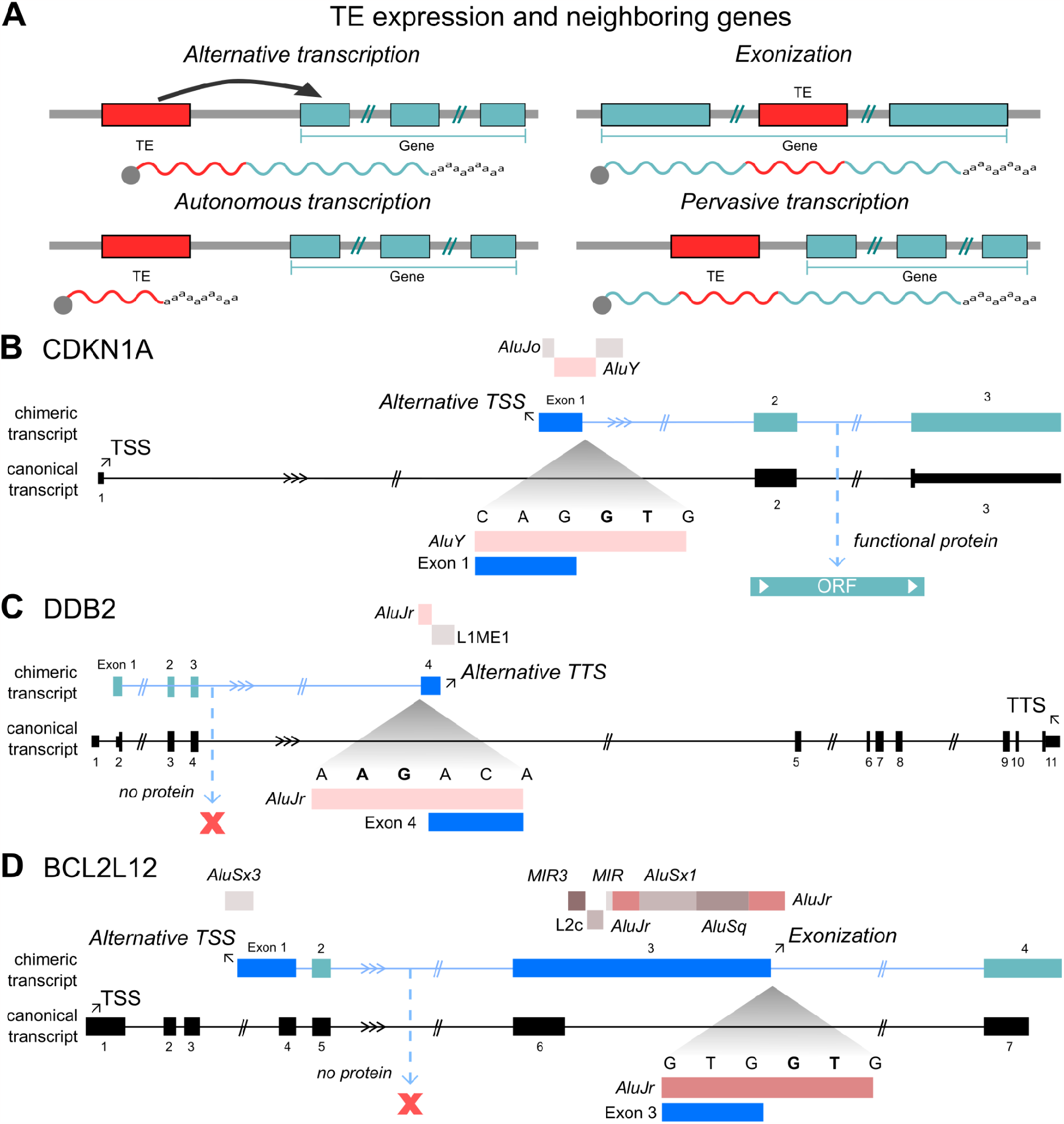
Transposable elements impact on host genes. **A)** Possible consequences of transposable element expression. **B)** *CDKN1* chimeric transcript showing an alternative start site using an *Alu* sequence and the retention of *CDKN1* ORF. **C)** *DDB2* chimeric transcript showing an alternative termination site in an *Alu* sequence. **D)** *BCL2L12* chimeric transcript showing multiple exonized TEs.

### Enrichment of TE-associated genes are linked to treatment response

Lastly, we examined the functional enrichment of DEGs that either form chimeric transcripts or have TEs located upstream or within their sequences, based on the GO analysis (**Fig. 6A-B**; **Supplementary Table 11**). Notably, in the A2780 cisplatin-sensitive cell line under treatment, genes linked with TEs were implicated in the negative regulation of transcription by RNA polymerase II, consistent with a cell undergoing apoptosis. In contrast, the A2780cis cell line exhibited enrichment in biological processes related to DNA conformation changes and nucleosome assembly. These processes have previously been associated with cancer cell resistance to DNA-damaging drug treatments^28^. Consequently, our findings validate that TEs correlate with gene pathways consistent with the predicted cellular responses to cisplatin treatment.

**Figure 6.**
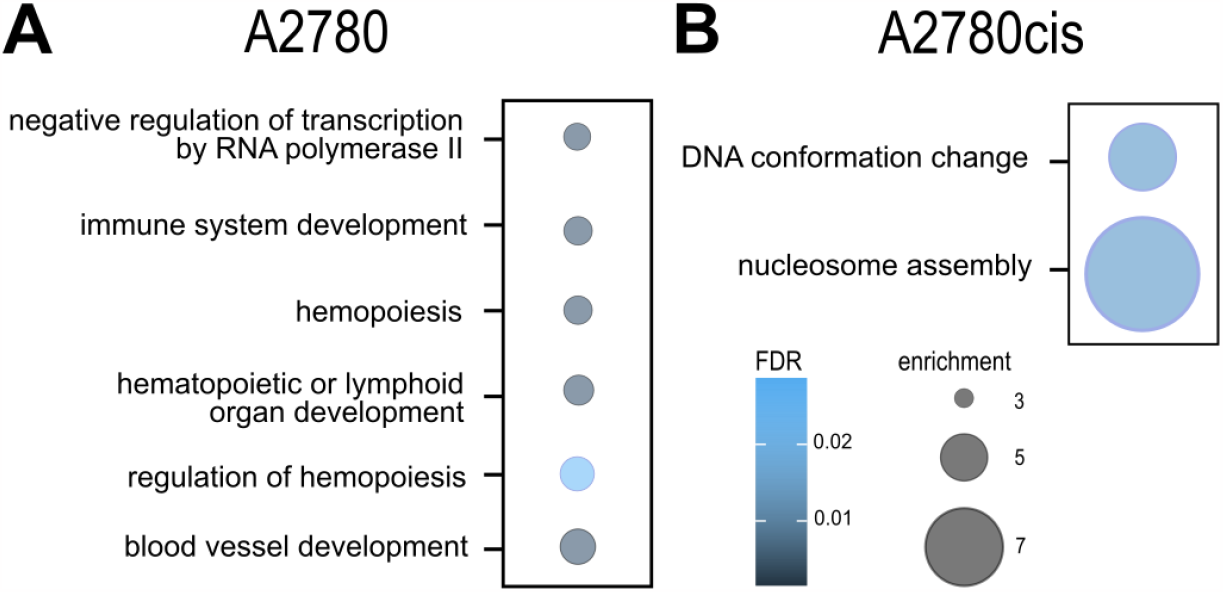
Biological processes for up-regulated DEGs including at least one gene with a TE either inside, upstream or participating in chimeric transcripts for. **A)** A2780, the cisplatin-sensitive cell line, and **B)** A2780cis, the cisplatin-resistant. Genes filtered for FDR below 5% and log 2-fold change higher than 1.

## DISCUSSION

Although it is widely recognized that TEs play a pivotal role in regulating gene expression, especially in complex diseases like cancer^10^, their roles in drug resistance mechanisms remain underexplored. In this study, we investigated the effects of cisplatin on both cisplatin-sensitive and -resistant OC cell lines, focusing on how TEs influence gene expression and are influenced by cisplatin treatment. We provided evidence that cisplatin treatment impacts TEs in addition to genes. Notably, our data revealed a distinct TE expression profile between cisplatin-sensitive and -resistant cell lines, as well as between treated and untreated cells.

The influence of TEs in the expression of genes is being reinforced by increasing evidence^10,14,16^. In examining this, we found that numerous DEGs had TEs situated either near or within their regulatory or transcribed domains. It remains unclear (and our data does not assert) whether such TEs are transcribed due to or as a result of these genes’ heightened expression. To provide clarity on this matter, we highlighted an expression pattern of sequences derived from ERVs in regulatory regions (attributing to the inherent transcriptional capability of such LTRs) and also pinpointed evidence of *Alu* and L1 expression stemming from the internal regions (introns) of DEGs, which are more prone to exonization.

Our investigations into chimeric transcripts of TEs and protein-coding genes yielded promising results. We discerned a collection of cancer-associated genes that formed chimeric sequences with TEs. Notably, we observed instances of exonization in genes that potentially amplify the treatment response without undermining gene functionality. In contrast, when exonization occurred in genes potentially promoting treatment evasion, the resultant chimeric transcripts were non-functional. We propose that, given their pervasive presence across the genome and their location within or near various genes’ introns, the activation of these elements might function synergistically. This could intensify the treatment response, as seen in our A2780 observations. However, under different conditions or treatments, these TEs could exert the opposite effect, fostering resistance to therapy.

In conclusion, we discovered that TEs, which are widespread throughout the genome, are altered collectively and appear to alter cancer-related gene transcription in response to cisplatin treatment. Our research not only provides findings in the field, but also vital insights that highlight the need of a holistic approach when examining cellular transcriptomes in complex models. To fully comprehend treatment response, we must extend our focus beyond protein-coding genes (or lncRNAs) and evaluate the entirety of all transcribed elements in our genome.

## METHODS

### RNA-seq data processing

We obtained RNA-seq and ATAC-seq data from the cisplatin-sensitive A2780 OC cell line and its resistant derivative, A2780cis, available in GenBank (NCBI) under accession GSE173201. The data includes the p53-proficient, cisplatin-sensitive A2780 OC cell line and cisplatin-resistant A2780cis, which was created after continuously exposing A2780 cells to increasing concentrations of cisplatin. We used RNA-seq and ATAC-seq data from A2780 control, A2780 treated with cisplatin, A2780cis control and A2780cis treated with cisplatin. For RNA-seq all samples were single-end libraries with three replicates per condition. For ATAC-seq all samples were paired-end data, with three replicates for controls and two for treatment. Cisplatin treatment consisted in 3 h of exposure to 200 µM of the drug. The dataset was downloaded from the NCBI Gene Expression Omnibus (GEO) database Sequence Read Archive using SRA toolkit (version 3.0.0; https://github.com/ncbi/sra-tools) and transformed to FASTq with fastq-dump. FASTQC^29^ software (version 0.11.9) was used to assess the sequencing quality of the data.

### Gene expression

Kallisto^20^ (version 0.48.0; --single -l 76 -s 1) was used to pseudo-align and quantify the gene expression data using the RNA-seq data (FASTq format) from A2780 and A2780cis using the release 36 of GENCODE^30^. Transcript level counts and transcripts-per-million (TPM) from Kallisto were aggregated to gene level counts and TPM using tximport^31^ R package (version 1.3.9; default parameters). Gene counts were used for differential expression analysis using DESeq2^25^ R package (version 1.34.0); genes with an adjusted p-value for multiple tests (False Discovery Rate (FDR)) < 0.05 and a log 2-fold change of at least -1 or 1 were considered differentially expressed. Genes annotated by the Catalogue Of Somatic Mutations In Cancer as hallmarks of cancer^32^ were used to identify tumor suppressor genes and oncogenes (version 94). Heatmaps were created using the clustermap function of the seaborn Python package (https://seaborn.pydata.org/), which employs average-link hierarchical clustering, and MA plots were created in Python using the bioinfokit^33^ toolkit.

### TE expression

REdiscoverTE^21^ software was used to gather TE expression data using the RNA-seq data (FASTq format) from A2780 and A2780cis. REdiscoverTE allows a whole-transcriptome RNA-Seq quantification simultaneously for TEs and protein-coding genes. Salmon^34^ (version 1.9.0; --seqBias, --gcBias, --validateMappings, --unmatedReads) was used to generate a quasi mapping index for REdiscoverTE repetitive elements raw counts. DESeq2 was used for differential expression analysis, and TEs with an adjusted p-value for multiple tests (FDR) < 0.05 and a log2 fold change of at least -1 or 1 were considered differentially expressed. The sequences classified as “low-complexity”, “satellite”, “simple-repeat”, “rRNA”, “tRNA”, “snRNA”, and “unknown” were excluded from the results.

### TE and host genes

To find genomic overlapping between DEGs and TEs, gene coordinates from GENCODE (release 36) and 5 kb upstream the beginning of each gene were intersected with TE coordinates (REdiscoverTE annotation) using BEDtools^24^ (version 2.27.1; intersect -s -f 0.2). Only DEGs with a log 2-fold change greater than 1 were used in this analysis. Biological processes with DEGs having TEs either inside or 5 kb upstream of their region were gathered.

To identify chimeric transcripts between TEs and protein coding genes, an internal pipeline (Mercuri RLV, Rangel A, and Galante PAF, in prep.) was used. Briefly, in this pipeline (Freddie) the RNAseq data from A2780 and A2780cis were aligned against the human reference genome (version GRCh38) using STAR^35^ (version 2.7.7a; --runThreadN 12), and only unique alignments were selected. Next, the aligned reads were assembled using StringTie2^36^ (version 2.1.4; -t 3) using GENCODE v36 and the human reference genome (GRCh38) as guide. To find the chimeric transcripts, we compared the newly assembled exons with the DFam^37^ intronic mobile elements annotations using BEDtools (intersect -r -f 0.5). To be considered a newly chimeric exon, at least 50% of the TE genomic region should be in this exon and 50% of this exon should be in the TE region. To assess whether the TE sequences found within chimeric transcripts were associated with alternative transcription start or termination sites, we used the UCSC BLAT^38^ tool followed by the Genome Browser^39^, and the results from RepeatMasker and RefSeq in human reference genome (GRCh38) were retrieved.

### ATAC-seq analysis of genes

Reads were aligned to the genome using Bowtie2^22^ (version 2.3.4.3; -very-sensitive -k 10). Following alignment, reads that were mapped to the mitochondrial chromosome were filtered and PCR duplicates were removed using PicardCommandLine MarkDuplicates (version 1.114). Insert size distribution histograms were plotted using deepTools^40^ (version 3.5.1; --maxFragmentLength 500) bamPEFragmentSize command. BAM files were converted to BED files in order to call for peaks. Peaks were called separately for each replicate BED file using MACS2^23^ callpeak command (version 2.2.8; --keep-dup all -g hs -f BEDPE)(Q-value (FDR) cut-off = 0.05). BEDtools was used to merge peak files from the same condition (intersect -r -f 0.3) and also to identify gained peaks of treatment compared to control (intersect -v -r -f 0.3). To find genomic overlapping between peaks and genes, gene coordinates from GENCODE (release 36) +1 kb upstream the beginning of each gene were intersected with peak coordinates using BEDtools (intersect -wo -f 0.3). Up-regulated and down-regulated DEGs with TEs either inside or upstream their sequences as detailed above (see TEs and host genes section) were retrieved.

### ATAC-seq analysis of TEs

Gained peaks of treatment were identified as described above. To determine the enrichment of peaks nearby each TE family in relation to the genomic abundance of such families (compared to a random shuffling of such TEs), we used the Perl script TE-analysis_Shuffle_bed.pl from the software TEanalysis (version 4.4.2; -l none -n 100 -o 10; https://github.com/4ureliek/TEanalysis) along with the TE BED file (REdiscoverTE annotation) + 1 kb upstream the beginning of each TE and 1 kb downstream the end of each TE. The significance of enrichment was estimated using binomial and two-tailed permutation tests. Results of up-regulated TEs (log 2-fold change higher than 0.5) either inside or upstream up-regulated DEGs in gained peaks were retrieved.

### Gene Ontology enrichment analysis of genes

A Gene Ontology (GO) analysis was performed using ShinyGO^26^ (version 0.76.2). First, for up-regulated DEGs found in ATAC-seq peaks with log 2-fold change greater than 1. Second, for DEGs with log 2-fold change greater than 1 and lower than -1, as separate sets, grouped into up-regulated and down-regulated. In order to better summarize and to remove redundancy, GO biological processes with more than two genes involved, FDR less than 0.05 and fold-enrichment greater than 2 were submitted to ReviGO^27^ (version 1.8.1; medium redundancy removal). Bubble plot was created using gapminder (https://www.gapminder.org/), ggplot2 and dplyr R packages (versions 3.4.0 and 1.0.10, respectively).

## Acknowledgements

The authors acknowledge partial support from CAPES and CNPq.

## Author Contributions

DMM was responsible for designing the study, conducting the search, extracting and analyzing data, interpreting results and updating reference lists. RLVM was responsible for analyzing part of the data. TMFFG provided feedback on the report. PAFG and ELSL contributed to the design of the study and results interpretation. DMM and PAFG were responsible for writing the manuscript with the input from all other authors.

## Data availability

The datasets analyzed during the current study are available in the GenBank (NCBI) under accession GSE173201.

## Notes

### Competing Interest Statement

The authors have declared no competing interest.

