## Supplementary Figures for "TRANSPOSABLE ELEMENTS ALTER GENE EXPRESSION AND MAY IMPACT RESPONSE TO CISPLATIN THERAPY IN OVARIAN CANCER"

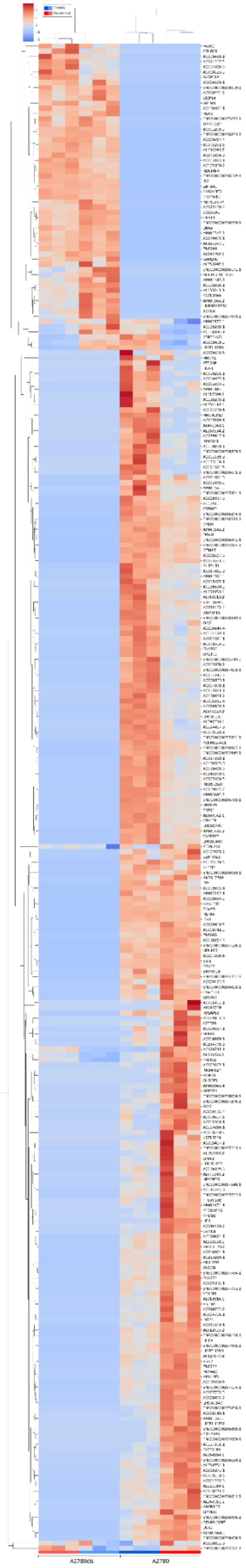


**Supplementary Figure 1. Heatmap of differentially expressed genes with TEs inside or upstream their sequences using Z-Score of TPM.** Genes were filtered adjusted P-value (FDR) below 5% and log 2-fold change lower than -1 or higher than 1.

Also provided as a separate PDF file.

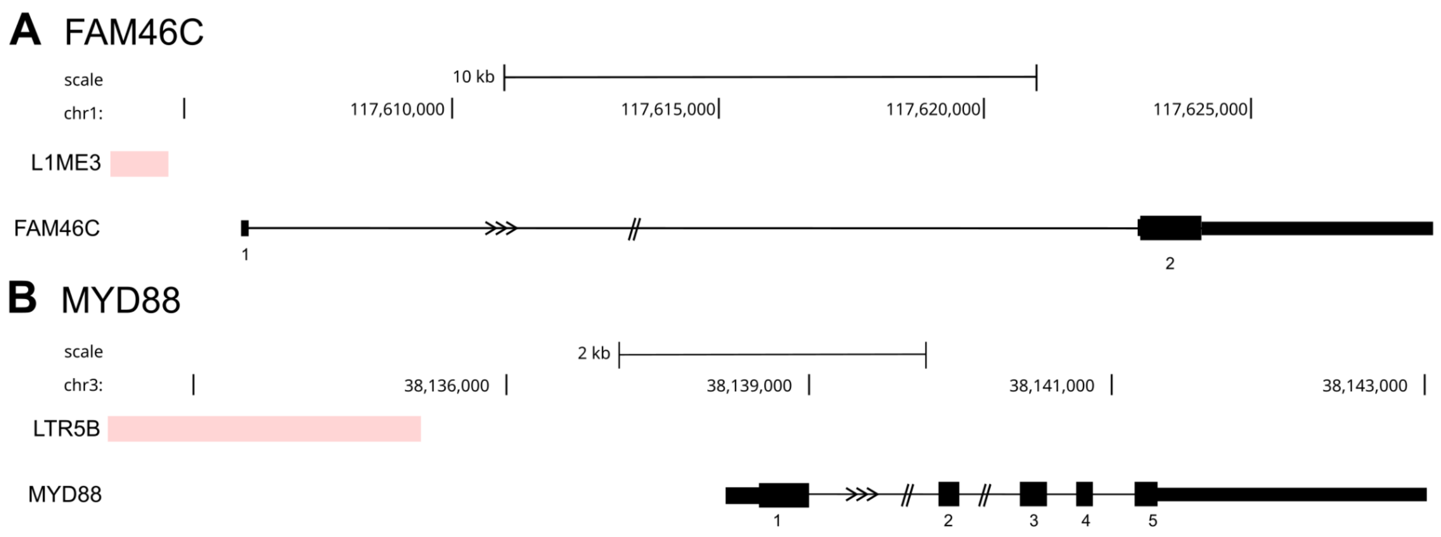


**Supplementary Figure 2. Up-regulated cancer-related genes and their associated TEs. A)** *FAM46C*, a tumor suppressor gene, with a L1ME3 upstream its sequence and, **B)** *MYD88*, an oncogene, with an ERVK (LTR5B) family member upstream its sequence.

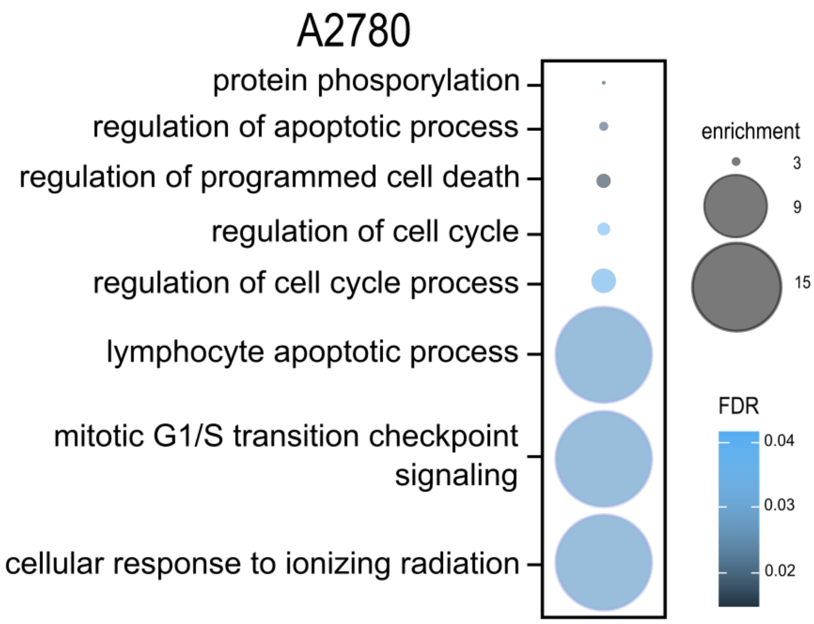


**Supplementary Figure 3. Biological processes for up-regulated DEGs found in peaks uniquely accessible after cisplatin treatment.** Genes filtered for FDR below 5% and log 2-fold change higher than 1. GO terms filtered for more than two genes involved, FDR below 5% and fold-enrichment score higher than 2.

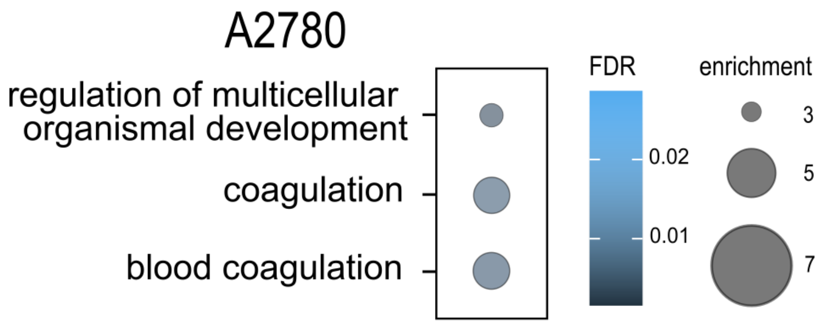


**Supplementary Figure 4. Biological processes for down-regulated DEGs including at least one with a TE either inside or upstream.** Genes filtered for FDR below 5% and log 2-fold change lower than -1. GO terms filtered for more than two genes involved, FDR below 5% and fold-enrichment score higher than 2.
