## Supplementary tables for "TRANSPOSABLE ELEMENTS ALTER GENE EXPRESSION AND MAY IMPACT RESPONSE TO CISPLATIN THERAPY IN OVARIAN CANCER"

**S2 Table. Differentially expressed protein-coding genes shared between A2780 and A2780cis cells.** Genes filtered for FDR below 5% and log 2-fold change higher than 1 or lower than -1.

| | A2780 | | | A2780cis | | |
|---|---|---|---|---|---|---|
| **Gene** | **log2FoldChange** | **p-value** | **adjusted p-value** | **log2FoldChange** | **p-value** | **adjusted p-value** |
| **ENSG00000284820.1** | 11.49470218 | 1.54E-261 | 5.91E-259 | 11.49470218 | 1.54E-261 | 5.91E-259 |
| **ENSG00000183793.14** | 10.02872507 | 4.15E-09 | 5.46E-08 | 10.02872507 | 4.15E-09 | 5.46E-08 |
| **ENSG00000030110.13** | 9.837973546 | 5.04E-09 | 6.55E-08 | 9.837973546 | 5.04E-09 | 6.55E-08 |
| **ENSG00000149346.15** | 9.176951079 | 0 | 0 | 9.176951079 | 0 | 0 |
| **ENSG00000223496.3** | 9.001746201 | 6.72E-08 | 7.16E-07 | 9.001746201 | 6.72E-08 | 7.16E-07 |
| **ENSG00000196866.3** | 8.690561729 | 0 | 0 | 8.690561729 | 0 | 0 |
| **ENSG00000270882.2** | 8.613856519 | 0 | 0 | 8.613856519 | 0 | 0 |
| **ENSG00000274997.2** | 8.583383474 | 0 | 0 | 8.583383474 | 0 | 0 |
| **ENSG00000274641.2** | 8.402270207 | 0 | 0 | 8.402270207 | 0 | 0 |
| **ENSG00000188396.4** | 3.296469766 | 5.02E-11 | 8.75E-10 | 3.296469766 | 5.02E-11 | 8.75E-10 |
| **ENSG00000120129.6** | 3.226247953 | 4.27E-110 | 1.18E-107 | 3.226247953 | 4.27E-110 | 1.18E-107 |
| **ENSG00000124216.4** | 3.176359562 | 5.45E-29 | 3.66E-27 | 3.176359562 | 5.45E-29 | 3.66E-27 |
| **ENSG00000185338.6** | 3.098689344 | 5.84E-26 | 3.40E-24 | 3.098689344 | 5.84E-26 | 3.40E-24 |
| **ENSG00000162772.17** | 2.943984864 | 6.07E-233 | 2.26E-230 | 2.943984864 | 6.07E-233 | 2.26E-230 |
| **ENSG00000181016.9** | 2.536636424 | 8.47E-08 | 8.85E-07 | 2.536636424 | 8.47E-08 | 8.85E-07 |
| **ENSG00000180245.7** | 2.312594863 | 3.38E-07 | 3.20E-06 | 2.312594863 | 3.38E-07 | 3.20E-06 |
| **ENSG00000130518.17** | 2.304499785 | 5.14E-14 | 1.25E-12 | 2.304499785 | 5.14E-14 | 1.25E-12 |
| **ENSG00000100652.5** | 1.99702803 | 3.11E-13 | 6.84E-12 | 1.99702803 | 3.11E-13 | 6.84E-12 |
| **ENSG00000196811.13** | 1.983799815 | 8.30E-08 | 8.69E-07 | 1.983799815 | 8.30E-08 | 8.69E-07 |
| **ENSG00000222009.8** | 1.876081469 | 3.85E-10 | 5.94E-09 | 1.876081469 | 3.85E-10 | 5.94E-09 |
| **ENSG00000131480.9** | 1.843598778 | 2.51E-10 | 3.95E-09 | 1.843598778 | 2.51E-10 | 3.95E-09 |
| **ENSG00000175906.5** | 1.74982994 | 1.33E-10 | 2.18E-09 | 1.74982994 | 1.33E-10 | 2.18E-09 |
| **ENSG00000130943.7** | 1.509758793 | 4.93E-07 | 4.54E-06 | 1.509758793 | 4.93E-07 | 4.54E-06 |
| **ENSG00000103269.14** | 1.446568763 | 5.78E-11 | 9.97E-10 | 1.446568763 | 5.78E-11 | 9.97E-10 |
| **ENSG00000172602.11** | 1.432879794 | 3.80E-07 | 3.57E-06 | 1.432879794 | 3.80E-07 | 3.57E-06 |
| **ENSG00000149243.16** | 1.426960768 | 1.32E-05 | 9.69E-05 | 1.426960768 | 1.32E-05 | 9.69E-05 |
| **ENSG00000187066.8** | 1.290509959 | 0.003422467469 | 0.01505470122 | 1.290509959 | 0.003422467469 | 0.01505470122 |
| **ENSG00000181773.7** | 1.24446698 | 3.37E-17 | 1.08E-15 | 1.24446698 | 3.37E-17 | 1.08E-15 |
| **ENSG00000186834.4** | 1.235145836 | 4.11E-108 | 1.12E-105 | 1.235145836 | 4.11E-108 | 1.12E-105 |
| **ENSG00000148926.10** | 1.217068503 | 3.12E-34 | 2.79E-32 | 1.217068503 | 3.12E-34 | 2.79E-32 |
| **ENSG00000126803.9** | 1.170140602 | 0.009519912595 | 0.03661138361 | 1.170140602 | 0.009519912595 | 0.03661138361 |
| **ENSG00000203811.1** | -1.031910863 | 7.41E-29 | 4.92E-27 | -1.031910863 | 7.41E-29 | 4.92E-27 |
| **ENSG00000126705.15** | -1.111728677 | 2.84E-17 | 9.20E-16 | -1.111728677 | 2.84E-17 | 9.20E-16 |
| **ENSG00000277075.2** | -1.314760222 | 4.40E-28 | 2.83E-26 | -1.314760222 | 4.40E-28 | 2.83E-26 |
| **ENSG00000184678.10** | -1.389580166 | 7.47E-133 | 2.23E-130 | -1.389580166 | 7.47E-133 | 2.23E-130 |
| **ENSG00000184357.5** | -1.419866519 | 3.58E-133 | 1.08E-130 | -1.419866519 | 3.58E-133 | 1.08E-130 |
| **ENSG00000187837.4** | -1.620384126 | 4.53E-132 | 1.34E-129 | -1.620384126 | 4.53E-132 | 1.34E-129 |
| **ENSG00000255423.1** | -2.755810464 | 7.26E-10 | 1.08E-08 | -2.755810464 | 7.26E-10 | 1.08E-08 |
| **ENSG00000124575.7** | -3.006222981 | 0 | 0 | -3.006222981 | 0 | 0 |
| **ENSG00000126860.12** | -3.345632646 | 0.0002726229397 | 0.00157841213 | -3.345632646 | 0.0002726229397 | 0.00157841213 |
| **ENSG00000285723.1** | -9.219486546 | 7.88E-08 | 8.28E-07 | -9.219486546 | 7.88E-08 | 8.28E-07 |

| | | | | | | |
|---|---|---|---|---|---|---|
| **ENSG00000170881.5** | -9.461547142 | 1.45E-08 | 1.73E-07 | -9.461547142 | 1.45E-08 | 1.73E-07 |
| **ENSG00000143751.10** | -9.469781221 | 0 | 0 | -9.469781221 | 0 | 0 |
| **ENSG00000013016.16** | -9.671855987 | 1.09E-08 | 1.34E-07 | -9.671855987 | 1.09E-08 | 1.34E-07 |

**S4 Table. Differentially expressed transposable elements shared between A2780 and A2780cis cells.** TEs filtered for FDR below 5% and log 2-fold change higher than 1 or lower than -1.

| | **A2780** | | | **A2780cis** | | |
|---|---|---|---|---|---|---|
| **TE** | **log 2-fold change** | **p-value** | **adjusted p-value** | **log 2-fold change** | **p-value** | **adjusted p-value** |
| **LTR27B** | 8.165620794 | 0.000001183664243 | 0.00001509490099 | 9.631372513 | 0.000000153893318 | 0.000003304793795 |
| **PrimLTR79** | -10.64553317 | 8.50E-85 | 8.40E-83 | 9.479468624 | 0.0000002722006693 | 0.000005653741771 |
| **X2_LINE** | -1.976165685 | 0.002784157391 | 0.01814866652 | -1.260989949 | 0.00002561355685 | 0.0004570757258 |

**S7 Table. GO enrichment analysis of up-regulated DEGs found in peaks uniquely accessible after cisplatin treatment.** Genes filtered for FDR below 5% and log 2-fold change higher than 1. GO terms filtered for more than two genes involved, FDR below 5% and fold-enrichment score higher than 2.

| **A2780 cisplatin-sensitive** | | | | | | | | |
|---|---|---|---|---|---|---|---|---|
| **ShinyGO** | **Up-regulated** | | **ReviGO** | **Up-regulated** | | | | |
| **TermID** | **Enrichment FDR** | **Fold Enrichment** | **TermID** | **Name** | **LogSize** | **Frequency** | **Uniqueness** | **Dispensability** |
| GO:0071479 | 0.03469275719 | 15.7976438 | GO:0006468 | protein phosphorylation | 6.115633474 | 4.340892012 | 0.993503702 | 0 |
| GO:0044819 | 0.03469275719 | 15.58700855 | GO:0010564 | regulation of cell cycle process | 4.97163761 | 0.3115866308 | 0.6583327806 | 0.22027307 |
| GO:0070227 | 0.03469275719 | 15.18215118 | GO:0042981 | regulation of apoptotic process | 4.996165675 | 0.3296910324 | 0.7770930124 | 0 |
| GO:0010942 | 0.03469275719 | 4.116287468 | GO:0043067 | regulation of programmed cell death | 5.012567288 | 0.3423804118 | 0.7766177019 | 0.2220954 |
| GO:0010564 | 0.04108875223 | 3.616221049 | GO:0044819 | mitotic G1/S transition checkpoint signaling | 2.959518377 | 0.00302682445 | 0.6559077414 | 0.65028213 |
| GO:0051726 | 0.04108875223 | 3.142542046 | GO:0051726 | regulation of cell cycle | 5.182183462 | 0.5059719579 | 0.7716889902 | 0.23077234 |
| GO:0043067 | 0.01202883398 | 3.12388087 | GO:0070227 | lymphocyte apoptotic process | 3.135768515 | 0.004543562856 | 0.99503425 | 0.00744137 |
| GO:0042981 | 0.01754590607 | 3.031335555 | GO:0071479 | cellular response to ionizing radiation | 3.608846822 | 0.01351094606 | 0.9419196645 | 0.00811455 |
| GO:0010941 | 0.01202883398 | 3.00642761 | | | | | | |
| GO:0006468 | 0.01754590607 | 2.960881391 | | | | | | |

**S8 Table. Results of the intersection between up-regulated DEGs with TEs inside or upstream and ATAC-seq peaks.** Genes filtered for FDR below 5% and log 2-fold change higher than 1.

| **A2780 cisplatin-sensitive** | | | | | | | | | | | | | | |
|---|---|---|---|---|---|---|---|---|---|---|---|---|---|---|
| **genes versus peaks from Suppl Table S6** | | | | | | | | | **genes versus TEs from Suppl Table S5** | | | | | |
| **chr** | **start** | **end** | **peak** | **chr** | **start** | **end** | **gene** | **log 2-fold change** | **chr** | **start** | **end** | **TE** | **TE subfamily** | **log 2-fold change** |
| chr10 | 43137382 | 43138393 | **peak_3651** | chr10 | 43136824 | 43138334 | **ENSG00000273008.1** | 1.315733343 | chr10 | 43137147 | 43137459 | **AluJb** | Alu | na |
| chr10 | 43138698 | 43139191 | **peak_3652** | chr10 | 43136824 | 43139334 | **ENSG00000273008.1** | 1.315733343 | chr10 | 43137147 | 43137459 | **AluJb** | Alu | na |
| chr13 | 111154066 | 111154398 | **peak_8330** | chr13 | 111148196 | 111154171 | **ENSG00000285856.1** | 2.150766136 | chr13 | 111149436 | 111152075 | **Tigger17** | TcMar-Tigger | na |
| chr14 | 34461308 | 34462688 | **peak_8588** | chr14 | 33924227 | 34462774 | **ENSG00000129521.15** | 1.370225167 | chr14 | 34464981 | 34466329 | **L2c** | L2 | na |
| chr18 | 14236472 | 14236857 | **peak_13928** | chr18 | 14225224 | 14342505 | **ENSG00000283294.1** | 9.265244459 | chr18 | 14223572 | 14225651 | **SVA_A** | SVA | 0.6848179153 |
| chr2 | 3575792 | 3576224 | **peak_16421** | chr2 | 3568505 | 3576100 | **ENSG00000287126.1** | 9.568299823 | chr2 | 3568504 | 3570040 | **SVA_D** | SVA | 0.5204788585 |
| chr2 | 87348356 | 87348684 | **peak_17192** | chr2 | 87311460 | 87348728 | **ENSG00000287763.1** | 1.320384817 | chr2 | 87349945 | 87351231 | **L1M1** | L1 | na |
| chr21 | 39313611 | 39313902 | **peak_19632** | chr21 | 39312935 | 39314962 | **ENSG00000255568.3** | 1.149265249 | chr21 | 39314756 | 39314988 | **MIR** | MIR | na |
| chr21 | 39314041 | 39314960 | **peak_19633** | chr21 | 39313935 | 39314962 | **ENSG00000255568.3** | 1.149265249 | chr21 | 39314756 | 39314988 | **MIR** | MIR | na |
| chr4 | 6987435 | 6987767 | **peak_22373** | chr4 | 6985926 | 6988036 | **ENSG00000287104.1** | 1.622777239 | chr4 | 6985872 | 6986478 | **L1ME1** | L1 | -0.5264188564 |
| chr5 | 72956553 | 72956941 | **peak_23846** | chr5 | 72955206 | 72956699 | **ENSG00000272081.1** | 1.859895614 | chr5 | 72957206 | 72958721 | **L1MB7** | L1 | na |

| | | | | | | | | | | | | | | |
|---|---|---|---|---|---|---|---|---|---|---|---|---|---|---|
| chr5 | 111223992 | 111224284 | **peak_24052** | chr5 | 111223653 | 111494886 | **ENSG00000152495.11** | 2.256534093 | chr5 | 111218019 | 111220205 | **L1MA4** | L1 | -0.8061290487 |
| chr5 | 111223992 | 111224284 | **peak_24052** | chr5 | 111223653 | 111494886 | **ENSG00000152495.11** | 2.256534093 | chr5 | 111221213 | 111222266 | **Tigger3b** | TcMar-Tigger | na |
| chr8 | 102806099 | 102806639 | **peak_28751** | chr8 | 102805517 | 102810039 | **ENSG00000253669.4** | 1.059486534 | chr8 | 102802260 | 102803410 | **L1MA3** | L1 | na |

**A2780cis cisplatin-resistant**

| **genes versus peaks from Suppl Table S6** | | | | | | | | | **genes versus TEs from Suppl Table S5** | | | | | |
|---|---|---|---|---|---|---|---|---|---|---|---|---|---|---|
| **chr** | **start** | **end** | **peak** | **chr** | **start** | **end** | **gene** | **log 2-fold change** | **chr** | **start** | **end** | **TE** | **TE subfamily** | **log 2-fold change** |
| chr1 | 34984903 | 34985676 | **peak_674** | chr1 | 34974356 | 34985313 | **ENSG00000284773.1** | 2.006290756 | chr1 | 34987583 | 34988922 | **Charlie10** | hAT-Charlie | na |
| chr10 | 71963950 | 71964385 | **peak_3370** | chr10 | 71963395 | 72013558 | **ENSG00000122863.6** | 8.915303256 | chr10 | 71961542 | 71962659 | **HERVFH21-int** | ERV1 | na |
| chr10 | 71963950 | 71964385 | **peak_3370** | chr10 | 71963395 | 72013558 | **ENSG00000122863.6** | 8.915303256 | chr10 | 71957210 | 71961280 | **HERVK14-int** | ERVK | na |
| chr10 | 71964865 | 71965398 | **peak_3371** | chr10 | 71964395 | 72013558 | **ENSG00000122863.6** | 8.915303256 | chr10 | 71961542 | 71962659 | **HERVFH21-int** | ERV1 | na |
| chr10 | 71964865 | 71965398 | **peak_3371** | chr10 | 71964395 | 72013558 | **ENSG00000122863.6** | 8.915303256 | chr10 | 71957210 | 71961280 | **HERVK14-int** | ERVK | na |
| chr22 | 22297467 | 22297896 | **peak_17011** | chr22 | 22297124 | 22329122 | **ENSG00000286129.1** | 9.770831971 | chr22 | 22295956 | 22297249 | **L1M1** | L1 | na |
| chr22 | 22298150 | 22298524 | **peak_17012** | chr22 | 22298124 | 22329122 | **ENSG00000286129.1** | 9.770831971 | chr22 | 22295956 | 22297249 | **L1M1** | L1 | na |

**S9 Table. TE enrichment in ATAC-seq peaks. TEs filtered for FDR below 5% and log 2-fold change higher than 0.5.** Binomial test and 2-tailed permutation test were filtered for P-values below 5%.

**A2780 cisplatin-sensitive**

| **Rclass** | **Rfam** | **Rname** | **obs_hits** | **2-tailed_permutation-test_pvalue(obs.vs.exp)** | **binomial_test_pval** |
|---|---|---|---|---|---|
| Retroposon | SVA | tot | 23 | 0.04 | 0.005333 |

**S10 Table. Results of chimeric transcripts analysis.** Genes filtered for FDR below 5% and log 2-fold change higher than 0.5 or lower than -0.5.

**A2780 cisplatin-sensitive**

***Alu***

| **gene** | **gene symbol** | **log 2-fold change** | **adjusted p-value** |
|---|---|---|---|
| ENSG00000168116 | KIAA1586 | 9.338047814 | 3.537041e-07 |
| ENSG00000154380 | ENAH | 8.832721048 | 1.285003e-06 |
| ENSG00000124762 | CDKN1A | 1.43697599 | 3.372629e-221 |
| ENSG00000134574 | DDB2 | 0.7115154913 | 1.494225e-13 |
| ENSG00000126453 | BCL2L12 | 0.5022478297 | 3.474373e-11 |
| ENSG00000096093 | EFHC1 | -0.5164685626 | 1.515155e-04 |
| ENSG00000073614 | KDM5A | -0.536998505 | 5.276004e-19 |
| ENSG00000122008 | POLK | -0.5457484 | 2.950297e-06 |
| ENSG00000086200 | IPO11 | -0.5591895889 | 5.916817e-07 |
| ENSG00000172530 | BANP | -0.5638836947 | 1.655231e-04 |
| ENSG00000068796 | KIF2A | -0.588912654 | 2.740053e-12 |
| ENSG00000078747 | ITCH | -0.6823700266 | 1.471804e-19 |
| ENSG00000153956 | CACNA2D1 | -0.6980388966 | 1.678698e-18 |
| ENSG00000108582 | CPD | -0.7369310935 | 1.740370e-28 |
| ENSG00000173611 | SCAI | -0.8006696836 | 4.291501e-11 |
| ENSG00000197056 | ZMYM1 | -0.9434494225 | 1.178340e-16 |

**L1**

| **gene** | **gene symbol** | **log 2-fold change** | **adjusted p-value** |
|---|---|---|---|
| ENSG00000166960 | CCDC178 | -0.6042673102 | 4.851764e-06 |

**A2780cis cisplatin-resistant**

**L1**

| **gene** | **gene symbol** | **log 2-fold change** | **adjusted p-value** |
|---|---|---|---|
| ENSG00000163704 | PRRT3 | 9.228712724 | 7.167677e-07 |

**S11 Table. GO enrichment analysis of DEGs with TEs forming chimeras, and upstream or inside its sequences.** DEGs were entered as two separate sets, up-regulated and down-regulated with log 2-fold change higher than 1 and lower than -1, respectively. GO terms filtered for more than two genes involved, FDR below 5% and fold-enrichment score higher than 2.

| **A2780 cisplatin-sensitive** | | | | | | | | |
|---|---|---|---|---|---|---|---|---|
| **ShinyGO** | **Up-regulated** | | **ReviGO** | **Up-regulated** | | | | |
| **TermID** | **Enrichment FDR** | **Fold Enrichment** | **TermID** | **Name** | **LogSize** | **Frequency** | **Uniqueness** | **Dispensability** |
| GO:0000122 | 0.0104513399577829 | 2.01014950905828 | GO:0000122 | negative regulation of transcription by RNA polymerase II | 4.71043894563884 | 0.170212441452898 | 0.895961839927603 | 0 |
| GO:0001568 | 0.00102952028785647 | 2.48177768362081 | GO:0001568 | blood vessel development | 4.53159386213364 | 0.112756583145298 | 0.328716481954613 | 0 |
| GO:0002520 | 0.00215426392411153 | 2.04469060779675 | GO:0002520 | immune system development | 3.937668314399 | 0.0287196401789159 | 0.445308837966418 | 0.57873533 |
| GO:0030097 | 0.00378213939569255 | 2.05245818446717 | GO:0030097 | hemopoiesis | 4.60763728141482 | 0.134334442563952 | 0.467235455309575 | 0.55027169 |
| GO:0048534 | 0.00138603878893057 | 2.11446479812751 | GO:0048534 | hematopoietic or lymphoid organ development | 3.72181061521255 | 0.0174698434660249 | 0.434173231473346 | 0.60677894 |
| GO:1903706 | 0.0276369992342764 | 2.39942520845346 | GO:1903706 | regulation of hemopoiesis | 4.25892448287688 | 0.0601812732495384 | 0.901017708563469 | 0.1810833 |
| **ShinyGO** | **Down-regulated** | | **ReviGO** | **Down-regulated** | | | | |
| **TermID** | **Enrichment FDR** | **Fold Enrichment** | **TermID** | **Name** | **LogSize** | **Frequency** | **Uniqueness** | **Dispensability** |
| GO:2000026 | 0.000723495902453451 | 2.30582880262991 | GO:2000026 | regulation of multicellular organismal development | 4.91612217879917 | 0.273323973422894 | 0.793799947586912 | 0.6287295 |
| GO:0050817 | 0.00535752002520516 | 3.56358826358826 | GO:0050817 | coagulation | 4.36466354089477 | 0.0767724853778338 | 0.983646818693534 | 0.49261771 |
| GO:0007596 | 0.00498581774470217 | 3.61135754326103 | GO:0007596 | blood coagulation | 4.36263324527236 | 0.0764144016628347 | 0.797145103610627 | 0.13815805 |

| **A2780cis cisplatin-resistant** | | | | | | | | |
|---|---|---|---|---|---|---|---|---|
| **ShinyGO** | **Up-regulated** | | **ReviGO** | **Up-regulated** | | | | |
| **TermID** | **Enrichment FDR** | **Fold Enrichment** | **TermID** | **Name** | **LogSize** | **Frequency** | **Uniqueness** | **Dispensability** |
| GO:0006334 | 0.0223029695353264 | 7.90567019247442 | GO:0006334 | nucleosome assembly | 4.43501574193256 | 0.0902735676692845 | 0.193922294111028 | 0.39664811 |
| GO:0071103 | 0.0223029695353264 | 4.69264166479527 | GO:0071103 | DNA conformation change | 5.37641634788993 | 0.7888153214738 | 0.327340939330734 | 0 |
